# Genome Assembly of *Astatotilapia latifasciata* Uncovers B Chromosome Linked Chromatin Reorganization

**DOI:** 10.1101/2025.08.05.668687

**Authors:** Maryam Jehangir, Syed Farhan Ahmad, Jordana Inácio Nascimento Oliveira, Adauto Lima Cardoso, Ivan Rodrigo Wolf, Guilherme Targino Valente, Kornsorn Srikulnath, Cesar Martins

**Author notes:** **Corresponding Author:** Cesar Martins Maryam Jehangir.

## Abstract

B chromosomes (Bs) are supernumerary genomic elements found in many eukaryotes, yet their full sequence composition, functional potential, and regulatory impact on the host genome remain unclear. Here, we present a chromosome-level genome assembly of the cichlid fish *Astatotilapia latifasciata*, integrating PacBio long reads, Illumina short reads, and Hi-C chromatin contact maps to resolve both A and B chromosomes. The 0.93 Gb assembly (N50 = 36.2 Mb) includes a 34 Mb B chromosome containing 789 predicted protein-coding genes and a markedly higher density of transposable elements (TEs), especially long terminal repeats (LTR) retrotransposons. Transcriptome profiling revealed that B-linked genes are predominantly transcriptionally repressed relative to their A chromosome paralogs. Hi-C based chromatin modeling uncovered distinct 3D structural configurations associated with the B chromosome, including fewer topologically associating domains (TADs), reduced loop formation, and altered compartmentalization. These changes are linked to long-range chromatin interactions and genomic rearrangements, suggesting that the B chromosome reshapes the nuclear architecture of the host genome. Our study proposes a potential regulatory role of Bs in genome and provides a genomic resource for investigating chromosome evolution in cichlids.

## Introduction

B chromosomes (Bs) are supernumerary, non-essential chromosomes present in a wide range of plant and animal taxa. Typically, heterochromatic and enriched in repetitive sequences, Bs are often considered selfish genetic elements that persist through non-Mendelian inheritance mechanisms such as meiotic drive (Camacho et al. 2000; Jones and Houben 2003; Houben 2017). Despite being dispensable for viability and development of the organism, the recent emergence role of Bs have been implicated in sex determination, and potential influence on A chromosomes gene expression, yet their precise evolutionary origin and functional impact remain unresolved (Camacho, 2005; Jones & Houben, 2003; Valente et al., 2017; Makunin et al., 2016). Current evolutionary models suggest that Bs originate from A chromosomes or from interspecific hybridization events, followed by amplification of repeats, segmental duplications, and structural rearrangements (Jones and Houben 2003; Houben et al. 2014; Valente et al. 2014; Ahmad and Martins, 2019). The maintenance of Bs often depends on drive mechanisms and escape from elimination during gametogenesis. Although Bs were traditionally viewed as inert genomic elements, increasing evidence points toward functional roles, including the expression of B-specific transcripts and modulation of host gene networks (Banaei-Moghaddam et al. 2013; Trifonov et al. 2013; Valente et al. 2014; Houben et al. 2019; Oliveira et al. 2024).

The cichlid fish *A. latifasciata* provides an emerging model to investigate B chromosome biology. This species harbors one or two morphologically identical Bs that coexist with 22 pairs of standard A chromosomes. Previous cytogenetic studies have identified ribosomal DNA clusters (Poletto et al. 2010), satellite DNA, and duplicated gene fragments on the B chromosome (Fantinatti et al. 2011; Carmello et al. 2017). Moreover, transcriptional comparative analysis between individuals with and without Bs (1B vs 0B) suggest potential regulatory impacts on gene expression (Ramos et al. 2017; Nascimento-Oliveira et al. 2021). While B chromosomes have been shown to harbor transcriptionally active sequences, the lack of a fully assembled B chromosome and the absence of Hi-C chromatin conformation data have limited efforts to define their regulatory landscape and spatial interactions with the A chromosome complement. Additionally, the high abundance of repetitive DNA on B chromosomes presents further limitations in characterizing their regulatory features and interactions. Hi-C sequencing has transformed our understanding of genome architecture, enabling the identification of topologically associating domains (TADs) and chromatin loops, at kilobase to megabase resolution (Lieberman-Aiden et al. 2009; Rao et al. 2015). TADs delineate functional domains that constrain enhancer-promoter interactions, and their disruption can impact transcriptional regulation. To date, no study has explored whether B chromosomes alter 3D genome topology or whether their structural features affect host nuclear organization. In this study, we generated the first chromosome-scale reference genome of *A. latifasciata* with a nearly assembled B chromosome, using deep coverage long-read sequencing and Hi-C chromatin contact maps. We assembled a 930 Mb genome, including a 34 Mb B chromosome, and performed transcriptomic, repeatome, and chromatin architecture analyses. Our results reveal that the B chromosome is transcriptionally less active, enriched in LTRs, and physically reorganizes chromatin interactions, supported from Hi-C comparative analysis. These findings provide new insights into the functional relevance of B and its evolutionary integration into host genome.

## Results

### Chromosome-Scale Assembly of the *A. latifasciata* Genome

We generated a chromosome-level reference genome for *A. latifasciata*, integrating PacBio long reads (∼118 Gb), Illumina short reads (∼51 Gb), and Hi-C chromatin interaction data (∼58 Gb) to produce a contiguous and accurate assembly of 930 Mb (**Table S1**). K-mer-based genome size estimation using Illumina data (1B male) predicted a genome of 902 Mb with low heterozygosity (0.49%) (**Table. S2**). PacBio scaffolds integrated with Hi-C mapping were anchored into 22 A chromosomes and a single B chromosome, covering 94% (872 Mb) of the genome and yielding an improved scaffold N50 of 36.2 Mb (**Table. S3**). Assembled chromosomes were ordered based on size (Chr1 to Chr22) (**Fig. 1a; Fig. S1a)** and substantially filled the gaps in the previous draft genome (**Fig. S1b**). The B chromosome was identified by comparing normalized read coverage between 1B (genome with B) and 0B (genome without B) individuals; scaffolds with high 1B/0B ratios were classified as B-linked. These were assembled into a 34.3 Mb long super-scaffold and validated using Hi-C interactions and known B-specific markers. Hi-C contact maps at high resolution (10–50 kb) revealed strong intra-chromosomal interactions in A chromosome, whereas Chr B displayed sparse interaction density, reflecting its isolated chromatin conformation (**Fig. 1a**). 3D genome in silico modeling showed Chr B localized toward the nuclear periphery with reduced chromatin compaction (smaller bead count), whereas A chromosomes (e.g., Chr 1) were centralized, consistent with active euchromatic regions (**Fig. 1b**). To benchmark assembly quality, we employed multiple metrics. K-mer spectra plots (**Fig. S2a**) revealed accurate haplotype resolution with minimal artifactual duplication. BUSCO analysis (*Actinopterygii* dataset, n = 3,640) recovered 96% complete orthologs (**Fig. S2b**), indicating high genome completeness. The LTR Assembly Index (LAI) averaged 8.5, with a maximum of 19.17, denoting high quality assembly for repetitive elements (**Fig. S2c; Table. S4**). The genome assembly was performed in a stepwise manner, progressing from initial draft versions (Asta_v1 and Asta_v2) to the final chromosome-scale assembly, Asta_v3. Compared to earlier drafts, Asta_v3 recovered approximately 150 Mb of previously missing or fragmented regions and improved assembly contiguity over 300-fold, increasing the scaffold N50 from 25 Kb to 36.4 Mb (**Fig. S2d, e; Fig. S1b; Table. S3**).

**Fig. 1.**
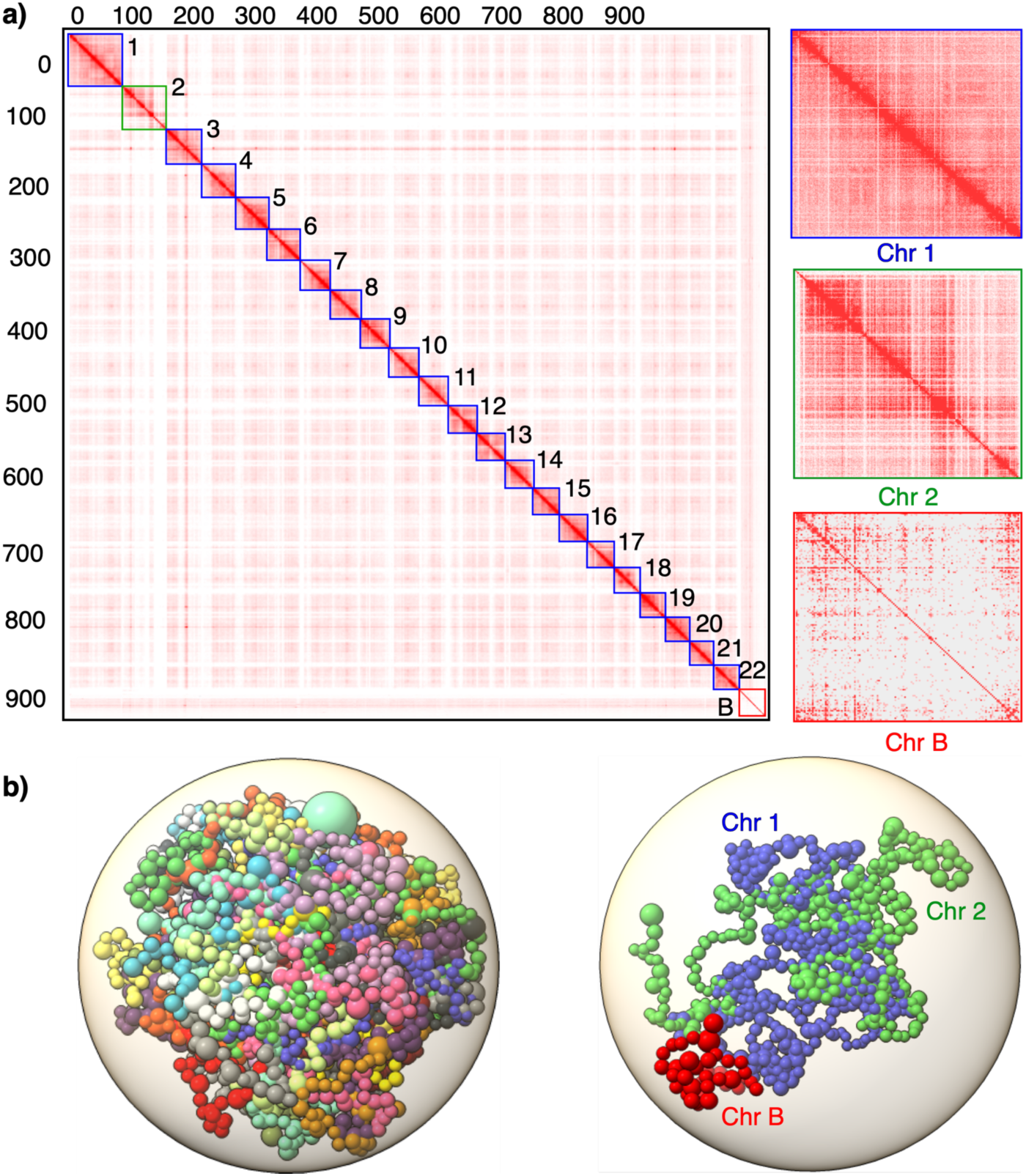
Chromosome-scale assembly and 3D genome architecture of *A. latifasciata* including the B chromosome. **(a)** Hi-C contact matrix (500 kb bins) showing chromatin interactions within and between chromosomes of *A. latifasciata*, including chromosomes 1–22 and the assembled “B_superscaffold” representing the B chromosome. Strong intrachromosomal interactions are evident along the diagonal, particularly between euchromatic arms of A chromosomes. Color intensity reflects the frequency of contact between pairs of 100-kb loci. White rows and columns represent bins with no valid interaction data. A magnified view (250 kb resolution) comparing the chromatin architecture of Chr 1, Chr 2, and Chr B is shown on the right. **(b)** Chrom3D structural model of the diploid genome (including the B chromosome). Each chromosome occupies a distinct territory within the nucleus (represented by different colors), with chromosomes modeled as continuous chains of beads, each bead corresponding to a predicted topologically associating domain (TAD). A separate 3D view highlights Chr 1, Chr 2, and Chr B in blue, green, and red, respectively.

### Genome annotation and repeatomics analysis revealed a low gene density and expansion of LTRs on B

Repeatome analysis revealed that approximately 39% of the *A. latifasciata* genome consists of repetitive DNA—an 11% increase over previous estimates based on short-read assemblies (Coan and Martins, 2016; Jehangir et al. 2019). This improvement reflects the enhanced resolution of complex repeats achieved through long-read sequencing. Most repeats were TEs, with DNA transposons comprising the largest fraction (23.1%). Among retroelements (13.4%), LINEs were the most abundant (∼6%), followed by LTR retrotransposons, with Gypsy elements accounting for 5.3% of the genome (**Table S5**). Insertion time analysis of LTRs indicated a relatively recent burst, beginning ∼300 Kya and peaking at a median age of 269 Kya (**Fig. S3; Table S6**). The genome also showed evidence of recent insertions from Gypsy, Copia, and unclassified LTR families (**Table S7**). Overall, LTR elements were distributed at a mean density of 10.45 per 10 Kb across the genome. Strikingly, Chr B exhibited a much higher LTR density, averaging 47.6 per 10 Kb—highlighting extensive LTR expansion on the B chromosome (**Fig. 2; Table S7**).

**Fig. 2.**
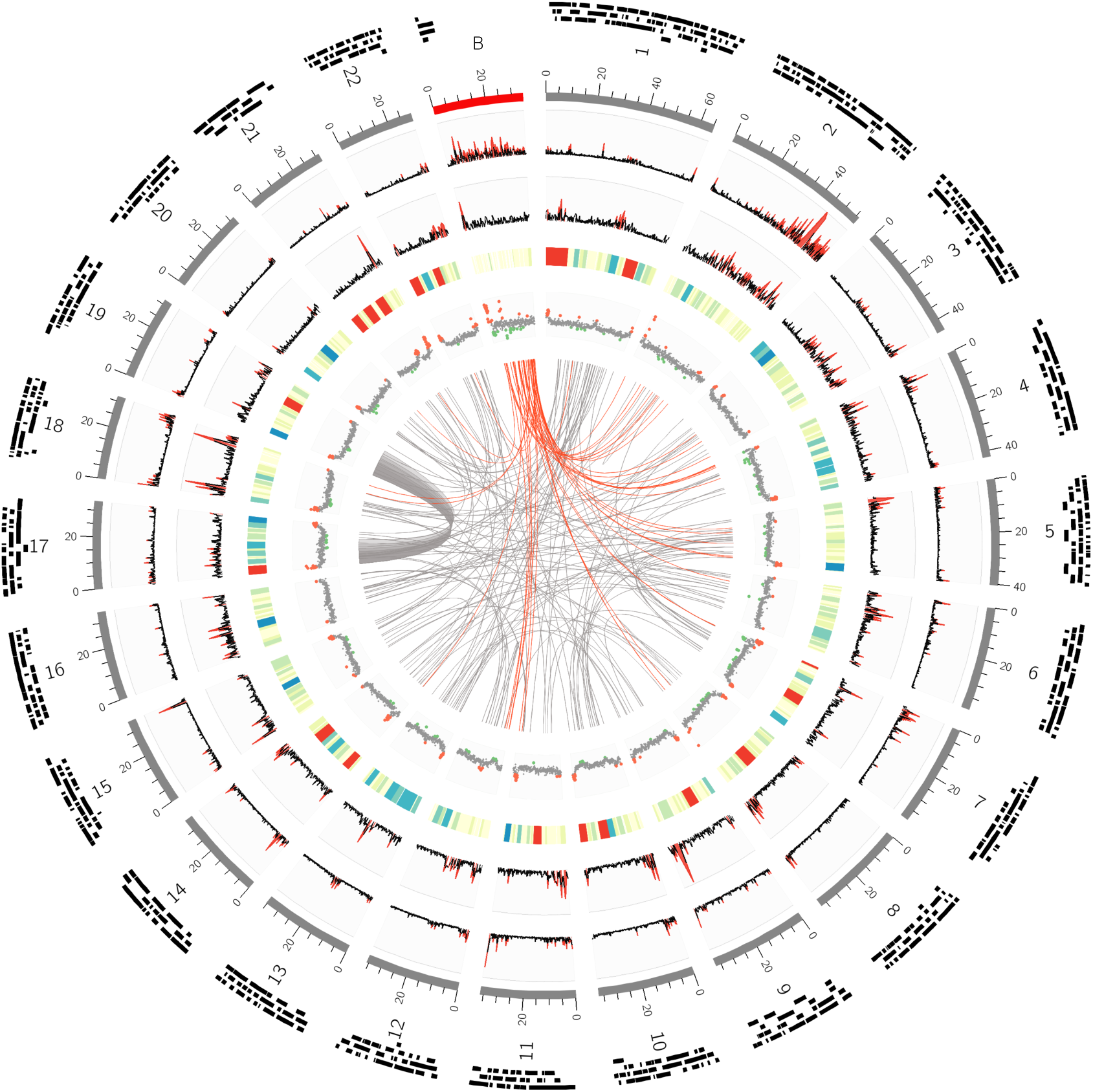
*A. latifasciata* genome assembly and chromosome-wide distribution of genomic features. The circos plot displays multiple layers of genome annotation and structure across assembled chromosomes, including the B_superscaffold. **Outermost ring:** Topologically associating domains (TADs) shown as black tiles. **Second ring:** Assembled chromosomes scaled by size (Mb), with A chromosomes in grey and the B_superscaffold in red. **Third and fourth rings:** Density of LTR retrotransposons and DNA transposons, respectively, calculated per 100 kb window. Red peaks denote regions with transposon enrichment (>30 TEs per 100 kb), notably concentrated on the B chromosome. **Fifth ring:** Gene density visualized as a heatmap—blue (30–60 genes/100 kb), red (>60 genes/100 kb) indicating gene-rich regions, and light yellow (<7 genes/100 kb) marking gene-poor regions, prominently observed on the B chromosome. **Sixth ring:** GC content displayed as a scatterplot (100 kb bins). Grey dots represent GC% between 40–45%, green dots <40%, and red dots >45%, with high GC variability particularly notable on the B chromosome. **Innermost links:** Syntenic duplications inferred from *SyRI* analysis. Red lines highlight duplicated regions involving the B chromosome.

Protein-coding genes were annotated on a repeat-masked version of Asta_v3 using an integrated approach that combined ab initio prediction with the *A. latifasciata* transcriptomes and proteomes from reference cichlid species. This resulted in 35,755 predicted protein-coding genes, with 33,146 (92.7%) assigned to the chromosomes, while remaining as unplaced scaffolds. Gene density was skewed toward smaller chromosomes, with Chr 21 and 22 being the most gene-rich (**Fig. 2; Table S7)**. The overall genome-wide mean gene density was 36.8 genes per 100 Kb, peaking on Chr 17 (85.7 genes/100 Kb) and lowest on Chr 12 (22.4 genes/100 Kb) (**Table S7**). Chr B had the lowest gene density at 7.9 genes per 100 Kb and carried 789 protein-coding genes in total (**Fig. 2**; **Table S7**). Additionally, GC content profiling in 10 Kb windows revealed highly variable patterns across Chr B, in contrast to the more uniform profiles observed on A chromosomes (**Fig. 2)**.

### An Integrative Genomic Approach Identified and Validated B Chromosome Sequences

While our chromosome-scale assembly resolved 22 A chromosomes, the identity of B chromosome scaffolds among unplaced sequences remained uncertain. To isolate and validate B-linked sequences, we implemented a three-tiered integrative strategy:

First, we performed a log₂ coverage ratio analysis using Illumina short-read data from 1B and 0B individuals, to recover scaffolds with significantly higher representation of mapped reads in 1B genome (**Fig. 3a**). Second, we applied a Hi-C–based comparative interaction frequency analysis between 1B and 0B contact maps. This identified a total of 283 unplaced scaffolds linked to B chromosome (**Table S8**) with significantly higher Hi-C interaction density in the 1B genome (**Fig. 3b, c**), consistent with their integration into a distinct chromosomal territory (**Fig. 1**). Furthermore, global coverage map of 0B and 1B genomic reads revealed B chromosome specific coverage peaks (highlighted as grey regions in **Fig. 3d**). Together, these two strategies allow us to recover a total of 34.4 Mb of B super-scaffold, placing the unplaced regions of the 1B assembly. Additionally, we mapped the remaining unplaced scaffolds against all assembled chromosomes, including the 22 A chromosomes and the B super-scaffold. This analysis revealed that most unplaced scaffolds shared homology with the B super-scaffold (**Fig. S4**), suggesting they are predominantly composed of repetitive sequences—a hallmark feature of the B chromosome. In the third approach, we validated B super-scaffold, by aligning previously characterized B-linked molecular markers—*ihhb* and BncDNA, previously experimentally confirmed and known for their localization on the B chromosome in *A. latifasciata* (Ramos et al. 2017; Jehangir et al. 2019). We observed the highest number of alignment hits for both *ihhb* (1,284 hits) and BncDNA (505 hits) on the assembled B super-scaffold (**Fig. 3e**). Other B-associated genes were also mapped to B super-scaffold, confirming the chr B assembly (**Table S9, S10**). Of particular interest, Chr 2 displayed several genomic mappings, and interaction features mirroring Chr B, including low gene density, high TE content, elevated mapping of B markers, and enrichment in unplaced scaffold alignments, suggesting a potential ancestral or structural relationship between Chr 2 and the B chromosome (**Fig. 3f, g**).

**Fig. 3.**
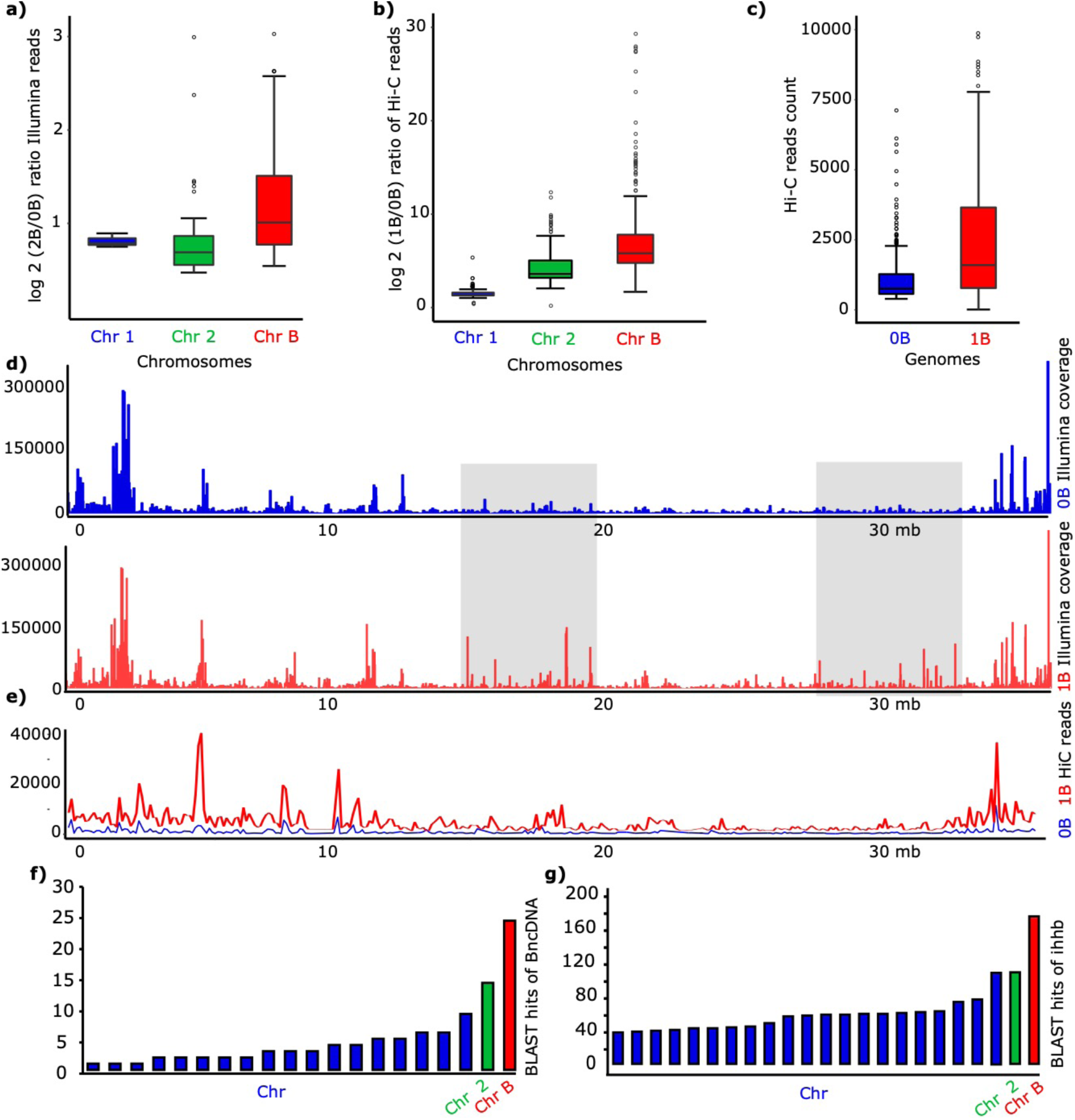
Validation of the B chromosome assembly. **(a, b)** Boxplots showing the distribution of log₂(2B/0B) Illumina read ratios and log₂(1B/0B) Hi-C read ratios across Chr 1, Chr 2, and Chr B, respectively, indicating B chromosome-specific enrichment. **(c)** Distribution of Hi-C reads mapped to the genome assemblies of 0B and 1B individuals. **(d)** Comparative coverage plot of Illumina reads mapped to the assembled B chromosome, with 0B (blue peaks) and 2B (red peaks) samples. Each peak represents read depth across 10 kb windows. Regions with substantial differences in coverage— lower in 0B and higher in 2B—are highlighted in grey. The X-axis indicates genomic position, and the Y-axis represents read depth. **(e)** Line plot comparing Hi-C interaction frequencies along the B chromosome between 0B (blue line) and 1B (red line) genomes, demonstrating increased interaction density in 1B. **(f, g)** Bar charts displaying the distribution of BLAST hits for B chromosome marker genes *BncDNA* and *ihhb*, confirming their enrichment on the B chromosome.

### TAD and Loop Architecture Are Altered in the 1B Genome

To investigate the impact of B chromosome presence on chromatin organization in host genome, we analyzed Hi-C data from both 0B and 1B individuals, mapping them to the chromosome-scale 1B genome assembly. We assessed chromatin contact profiles, TADs and loops, to compare 3D chromatin organization between the two genomes. Analysis of contact probability decay as a function of genomic distance revealed that both 0B and 1B genomes follow the expected power-law scaling, reflecting conserved chromatin folding principles. Notably, the 1B genome exhibited reduced short-range (<1 Mb) and increased long-range (>1 Mb) interactions compared to 0B, indicating a global reorganization of chromatin architecture. This was further supported by an elevated short-to-long (SVL) contact ratio across 50 kb bins, consistent with a genome-wide shift in chromatin compaction and folding dynamics potentially driven by the presence of B chromosomes (**Fig. 4a, b)**, which was also evident at the scale of individual chromosomes (Fig. S5).

**Fig. 4.**
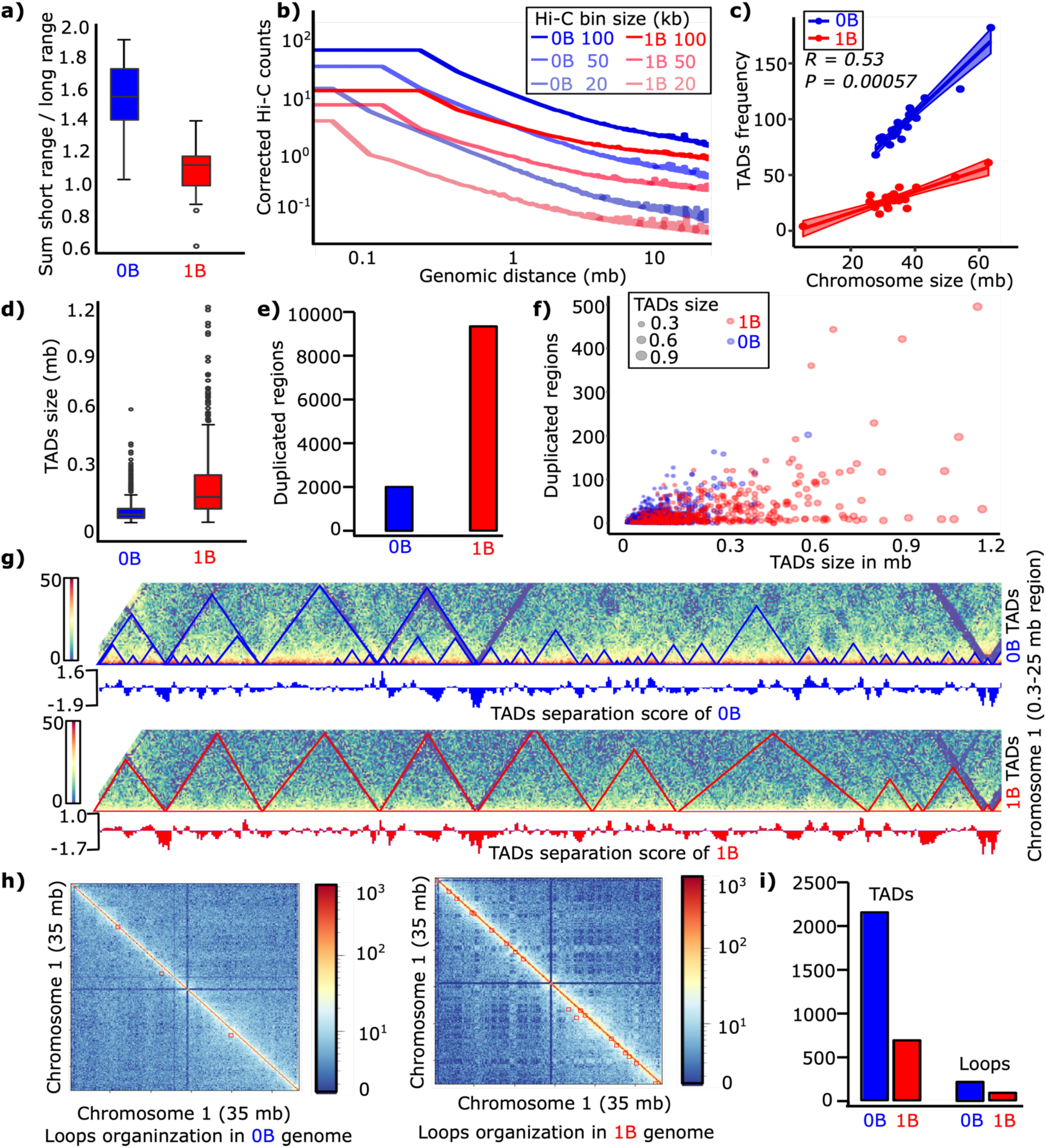
Comparative 3D genome architecture between 0B and 1B *A. latifasciata* genomes. **(a)** Boxplot comparison of short-range versus long-range chromatin contact frequencies in 0B and 1B genomes. **(b)** Hi-C contact decay curve showing contact frequency as a function of genomic distance at 50 kb resolution. The 0B genome (blue line) is enriched for short-range contacts (<1 Mb), while the 1B genome (red line) displays more long-range interactions (>1 Mb). **(c)** Correlation between the number of TADs (y-axis) and chromosome size (x-axis) in both 0B and 1B genomes. The 1B genome features longer but fewer TADs compared to the 0B genome. **(d)** Distribution of TAD sizes in 0B and 1B genomes. **(e)** Number of duplicated regions predicted by *SyRI* between the two genome types, and correlation between TAD size (x-axis) and the number of duplications (y-axis). Each point represents a TAD, colored by genome (blue = 0B, red = 1B) and scaled by TAD size. **(g)** High-resolution in situ Hi-C heatmaps of Chr 1 (25 Mb region) for 0B (top panel) and 1B (bottom panel) genomes at 50 kb resolution. TADs appear as “mountains,” with fewer, larger TADs in 1B. TAD separation scores (see Methods) are shown as tracks, with vertical lines indicating boundary positions. **(h)** Comparative Hi-C contact maps at 40 kb resolution showing loop structures (red dots) in 0B and 1B genomes. **(i)** Bar chart summarizing the total number of TADs and chromatin loops identified in each genome.

A total of 170 TADs were conserved between both genomes, the 1B genome exhibited fewer but significantly larger TADs (mean size 2.1 Mb vs. 0.7 Mb in 0B) (**Fig. 4 c, d**). We also observed a slight positive correlation (R = 0.53, p value < 0.001) between chromosome size and TAD frequency, indicating that larger chromosomes tend to harbor a greater number of TADs compared to smaller ones (**Fig. 4c**). Comparative analysis revealed a significant shift in TAD size distribution, with 1B exhibiting larger domains that span broader genomic regions but show reduced interaction density— indicative of weakened insulation or less well-defined TAD boundaries as reflected by a narrower and less negative insulation score range (**Fig. 4d**).

We then tested whether TAD reorganization in 1B was associated with structural rearrangements. Using SyRI (Goel et al. 2019), we identified 9,659 duplications in the 1B genome, and these duplicated regions overlapped significantly more with TAD boundaries than in 0B genome (**Fig. 4e, f**). This suggests that duplications may disrupt domain organization and contribute to TAD turnover. To further investigate the structural TADs organization, we examined TAD separation scores and found clear differences in domain insulation strength between 0B and 1B, with 1B exhibiting a narrower and less negative insulation score range (–1.7 to 1.0) compared to 0B (–1.9 to 1.6), indicative of weaker TAD boundaries (**Fig. 4g**). We identified 207 chromatin loops in 0B and only 67 in 1B, reinforcing the global reorganization in the 1B genome (**Fig. 4 h, i**). Additionally, comparison with Hi-C data from other cichlid species revealed a higher number of long-range interchromosomal interactions in 1B than in 0B or other genomes (**Fig. S6; Table S11**). These observations highlight that the B chromosome contributes to unique, large-scale chromatin reorganization in the host genome. These chromatin changes are also marked by highly rearranged 1B genome, with a total of 1,885 inversions (INV), 2,278 translocations (INVTR), and other SVs variations (**Fig. S7**). Comparison of B- and B+ genomes further identified that 1B genome has higher number of duplications as compared to 0B suggesting the B chromosome sequences might have duplicated at large scale during its evolution (**Table S12**). We also recognized multiple intra-chromosomal INVTRs on Chr 1, a large intra-chromosomal INV on Chr 2, as well as inter-chromosomal INVs between different chromosomes (**Fig. S7**).

### B Chromosome Genes Exhibits Transcriptional Repression

To evaluate the transcriptional expression of genes located on the B chromosome, we analyzed RNA-seq expression data from brain, muscle, and gonadal tissues of 1B and 0B individuals (both male and female). On average, genes located on Chr B exhibited markedly lower expression (mean logCPM = 1.84) compared to those on A chromosomes (mean logCPM > 4.58) across all tissues (**Fig. 5; Table S13**). When comparing expression levels across representative chromosomes (Chr 1, Chr 2, and Chr B), we observed consistently lower expression level (p value < 0.05) for Chr B genes, with Chr 2 also showing lower activity than Chr 1 (**Fig. 5 a-f**).

**Fig. 5.**
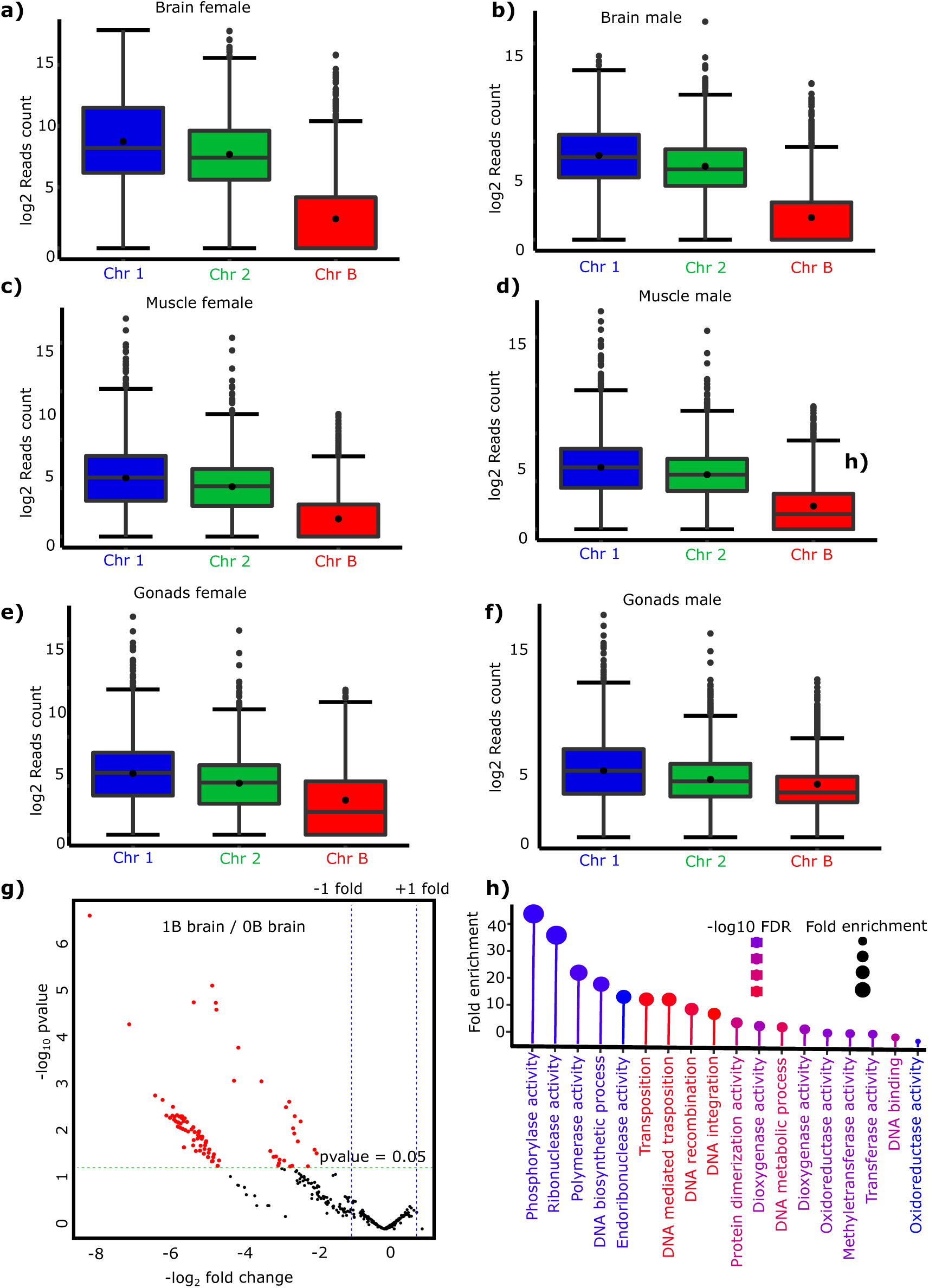
Transcriptional profiles of B chromosome genes reveal reduced expression relative to A chromosome genes. **(a–f)** Boxplots showing log₂-transformed RNA-seq read counts for genes located on Chr 1, Chr 2 (representative A chromosomes), and Chr B across various tissues from 1B individuals. In all cases, genes on the B chromosome exhibit significantly lower expression levels (p value < 0.05) compared to A chromosomes. Similar patterns were observed for the remaining autosomes (not shown). **(g)** Volcano plot of differentially expressed genes (DEGs) located on the B chromosome between 0B and 1B brain samples. The x-axis shows log₂ fold change (FC), and the y-axis shows –log₁₀(adjusted p-value). Genes significantly downregulated in 1B (padj < 0.001, log₂ FC < –2) are highlighted in dark red. Additional DEG results for gonad and muscle tissues are provided in the Supplementary Data. **(h)** Lollipop plot illustrating enriched functional categories of genes annotated on the B chromosome. Dot size corresponds to fold enrichment, and color intensity represents –log₁₀(FDR) values, providing a functional overview of the transcriptionally active regions on Chr B.

To further assess the regulatory impact of B-linked genes across different tissues, we performed differential expression analysis between 0B and 1B individuals using the edgeR package (Robinson et al., 2010). Consistent with the global repression of B chromosome genes, only a small number of differentially expressed genes (DEGs) were identified in muscle and gonadal tissues (**Fig. S8; Table S14**). This supports the interpretation that the B chromosome is largely transcriptionally inactive. If Chr B were broadly transcribed, we would expect widespread expression differences between 0B and 1B transcriptomes. Notably, a distinct pattern emerged in brain tissue, where several genes were significantly downregulated in 1B individuals (**Fig. 5g**, **Table S15**). This suggests that B chromosome gene expression may be cell type–specific, with functions related to hemostasis, spermatogenesis, skin pigmentation, defense response to hatching behavior and others. These findings suggest the selective transcriptional repression of B-linked genes and highlight tissue-specific regulatory effects associated with the presence of the B chromosome (Oliveira et al. 2024).

To explore the global functional repertoire of Chr B genes, we annotated all 789 protein-coding genes using Blast2GO, involving BLASTx homology search, GO term mapping, and annotation. Of these, 649 genes showed homology to known proteins, while 140 (17.7%) had no hits in current databases, indicating potential novel or B-specific genes. GO mapping assigned functional terms to 115 genes (19.2%), primarily sourced from the UniProtKB database. The top-hit species distribution showed *Oreochromis niloticus* as the closest homolog for most annotated genes. In the molecular function category, glycogen phosphorylase activity had the highest representation (88%), followed by RNA–DNA hybrid ribonuclease activity (**Table S16**). In the biological process category, GO enrichment highlighted functions related to DNA transposition (16%), DNA integration, and metabolic and recombination-related processes (**Fig. 5h**; **Table S16**). Together, these findings indicate that the B chromosome in *A. latifasciata* harbors largely transcriptionally silenced genes. However, the presence of a subset of active and potentially functional genes—some of which are potential novel—suggests that the B chromosome may behave as regulatory influence within the host genome.

### Comparative Genomics Reveals Evolutionary Dynamics and B Chromosome– Linked Gene Family Expansion

To contextualize the evolutionary history of *A. latifasciata* and assess the potential evolutionary adaptation signatures, we performed comparative phylogenomics and gene family evolution analyses across 16 teleost genomes, including 11 cichlids and 5 non-cichlid outgroups. A species tree constructed using 384 single-copy orthologs confirmed the monophyly of *A. latifasciata*, *Pundamilia nyererei*, *Metriaclima zebra*, and *Astatotilapia calliptera*. *A. latifasciata* and *P. nyererei* formed the most closely related pair, with a divergence time estimated at ∼4.83 Mya (**Fig. 6a; Fig. S9**). The Victoria and Malawi lake lineages diverged ∼6.14 Mya, while African and American cichlids split ∼65.6 Mya (**Fig. 6a**).

**Fig. 6.**
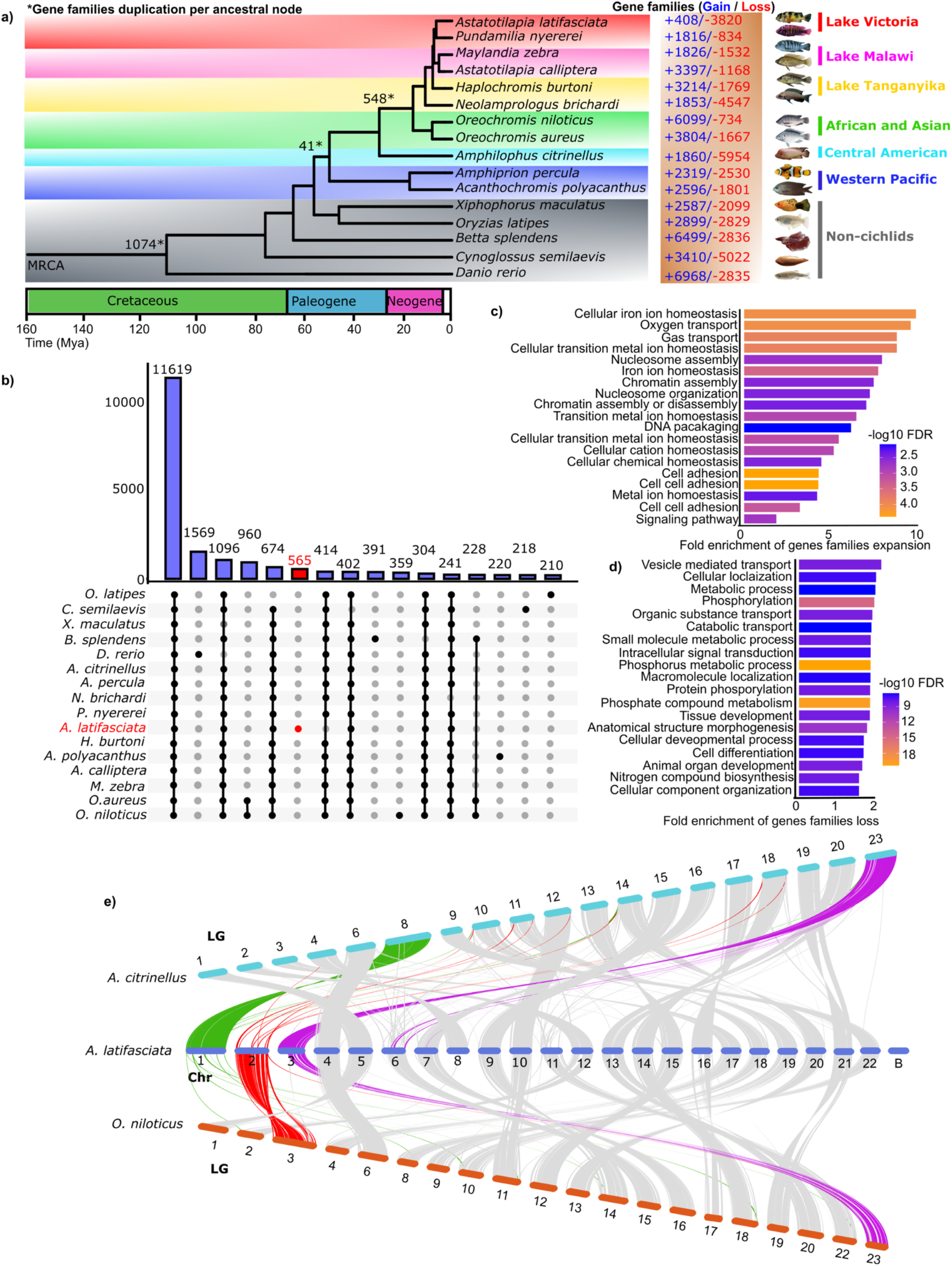
Evolutionary analysis of cichlid genomes and associated gene families, including *A. latifasciata*. **(a)** Time-calibrated phylogenetic tree of 11 representative cichlid species, with teleost outgroups. Gene family expansions (blue) and contractions (red), as identified by CAFE analysis, are indicated along branches and nodes. Numbers marked with asterisks denote predicted gene duplications inferred from gene clustering at each ancestral node. Divergence times are indicated with shaded boxes in units of million years ago (Ma). MRCA: most recent common ancestor. **(b)** UpSet plot showing the number of unique and shared gene families across species groups. *A. latifasciata* is highlighted in red. **(c, d)** Bar plots showing Gene Ontology (GO) enrichment of expanded (c) and contracted (d) gene families. The x-axis represents fold enrichment, while the y-axis lists GO terms, color-coded by –log₁₀(FDR) values. **(e)** Genome-wide collinearity among three cichlid species assemblies. Syntenic blocks highlight chromosomal rearrangements, with green and red regions on Chr 1 and Chr 2 of *A. latifasciata* representing fusion like events. Chr 3 is highlighted in purple and carries *amh*, homologous to the sex-determining LG23 in other cichlid species.

Orthogroup analysis clustered 97.1% of all genes into 30,304 gene families, including 4,443 species-specific orthogroups. *A. latifasciata* exhibited the highest number of unique gene families (n = 565), followed by *O. niloticus* (n = 359), suggesting substantial lineage-specific diversification (**Fig. 6b; Tables S17–S18**). A total of 29,679 genes from *A. latifasciata* were assigned to 16,666 gene families. Among these, 7,395 gene families were conserved and shared across all analyzed species, reflecting a core set of orthologous genes.

To model gene family dynamics, we used a stochastic birth–death process (see Materials and Methods) to infer lineage-specific gene expansions and contractions. In *A. latifasciata*, 3,820 gene families were contracted, while 408 expanded indicating a net gene family loss relative to related lineages such as Lake Victoria and Malawi cichlids (**Fig. 6a**). In contrast, *O. niloticus* showed substantial gene family expansion (n = 6,099) and minimal contraction (n = 734). At the teleost most recent common ancestor (MRCA), we identified 1,074 duplicated gene families.

Gene ontology enrichment of significantly expanded families (FDR < 0.05) in *A. latifasciata* revealed functional enrichment for genes involved in ion homeostasis, oxygen transport, and chromatin assembly (**Fig. 6c; Table S19**). These categories suggest possible roles in osmoregulatory adaptation and epigenetic remodeling. Conversely, contracted gene families were enriched in functions related to vesicle transport, catabolism, and developmental regulation (**Fig. 6d; Table S20**).

Remarkably, 16 of the rapidly expanded gene families were localized to the B chromosome (**Table S21**). Many of these genes lacked homology to reference databases, indicating possible novel B-linked genes. Others, such as *slc24a5*, *smg1*, *rgs2*, *atp5f1e*, and *histone1-like* genes, exhibited partial homology and were enriched in functions related to morphogenesis, chromatin organization, and DNA conformation (**Fig. S10; Table S22**). These chromatin and DNA related enriched functions further support the link between B chromosome–specific gene expansion and the large-scale 3D chromatin remodeling observed in 1B individuals (**Fig. 4**).

To explore chromosome evolution and structural rearrangements in cichlids genome, we performed interspecies synteny analyses using genome assemblies of *A. citrinellus* and *O. niloticus*. We observed high chromosomal synteny (>83%) between *A. latifasciata* and these reference genomes. Notably, Chr 2 of *A. latifasciata* appeared to result from ancestral fusion events involving multiple linkage groups (LGs) of *A. citrinellus*, while Chr 1 likely derived from a fusion of LGs present in *O. niloticus* (**Fig. 6e**). Chr 3 of *A. latifasciata* was syntenic with LG23 in both species, a region harboring the sex-determining gene *amh*. Additionally, our survey of known master sex-determining genes in cichlids revealed multiple significant matches on chromosome 3 (**Table S23**), suggesting that Chr3 harbors a putative sex determination region in *A. latifasciata*. Further comparison identified conserved scaffolds and ancestral rearrangement signatures, including chromosomal inversions and fusions, shared with *M. zebra* and *A. aureus* (**Fig. S11**). These syntenic relationships underscore the dynamic nature of cichlid genome evolution, shaped by lineage-specific fusions and duplications events.

## DISCUSSION

This study presents the chromosome-level genome assembly of an important cichlid species being extensively studies as model for chromosome biology and evolution, providing new insights into the structural, regulatory, and evolutionary roles of supernumerary chromosomes. Using a hybrid sequencing approach combining PacBio long reads, Hi-C chromatin conformation capture, and Illumina short reads, we assembled a 930 Mb reference genome for *A. latifasciata*, which includes 22 A chromosomes and a 34.3 Mb B chromosome. This high-contiguity assembly resolved ∼150 Mb of previously fragmented or missing regions, marking a significant improvement over earlier draft genomes (Jehangir et al. 2019) and enabling in-depth exploration of the B chromosome’s sequence content and biological impact. Inter-chromosomal homology analysis further revealed that many B-linked sequences had substantial homology with Chr 2. This supports a model in which the B chromosome originated primarily from Chr 2 through segmental duplications, transpositions, and rearrangements, with additional contributions from other A chromosomes, followed by sequence divergence and extensive TE accumulation (Camacho, 2005, Valente et al., 2014).

Our results support the view that B chromosomes are largely heterochromatic and enriched in repetitive sequences, particularly LTR retrotransposons, consistent with previous reports across diverse taxa (Jones and Rees, 1982; Camacho, 2005; Coan and Marins, 2018; Ramos et al., 2017; Ahmad et al., 2020). Although the B chromosome encodes 789 protein-coding genes, it displays markedly lower gene density and transcriptional activity compared to A chromosomes (mean logCPM = 1.84 vs. >4.58), suggesting widespread transcriptional repression. This likely reflects dosage regulation or epigenetic silencing mechanisms that limit the impact of B-linked sequences on host genome function. Notably, differential expression analysis revealed that B-linked genes were significantly expressed only in brain tissue, indicating that this repression is not uniform and may be relaxed in a tissue-specific manner. These findings highlight the potential for context-dependent regulatory activity of B chromosomes.

Hi-C analysis revealed that the presence of the B chromosome is associated with large-scale reorganization of 3D genome structure, particularly at the level of TADs. The 1B genome exhibited fewer but significantly larger TADs compared to the 0B genome, indicative of weakened domain insulation and altered chromatin compartmentalization. This shift in TAD organization was accompanied by an increase in long-range chromatin interactions and a decrease in short-range contacts, as quantified by SVL contact ratios. These results suggest that the B chromosome alters the folding landscape of the host genome, potentially through disruption of preexisting domain boundaries.

Notably, TADs in the 1B genome frequently overlapped with duplicated genomic regions, consistent with models in which structural variation drives TAD fusion or boundary erosion (Franke et al., 2016; Bonev and Cavalli, 2016; Szabo et al., 2019). Such architectural changes may have functional consequences by reshaping regulatory domains and altering the spatial organization of the genome. Moreover, the identification of species related expanded gene families on Chr B particularly those related to chromatin assembly, and DNA conformation, hints for B chromosome linked role in chromatin architecture modulation

From an evolutionary perspective, our orthogroup and gene family comparative analysis revealed a net loss of gene families in *A. latifasciata*, but also lineage-specific expansion of 408 families, including 16 localized on the B chromosome. These expanded gene families were enriched in homeostatic and oxygen transport functions, suggesting potential roles in species related environmental adaptation. Importantly, many of the B-linked expanded genes lacked homologs in current databases, indicating a reservoir of novel or rapidly evolving sequences. These results echo findings in other organisms where B chromosomes have been proposed for their potential role in genome innovation and adaptation (Miao et al. 1991; Ahmad et al. 2020; Johnson Pokorná and Reifová 2021; Liu et al. 2025).

While these findings collectively support the hypothesis that B chromosomes are not merely passive passengers but potentially dynamic contributors to genome architecture and evolution, our study has limitations. Despite the high contiguity of our assembly, there is cytological evidence that the B chromosome of *A. latifasciata* has a similar size to chromosome 1 (Poletto et al. 2010; Fantinatti et al. 2011; Cardoso et al. 2022), which indicates that the B chromosome was not fully resolved due to its extreme repeat content, ampliconic regions, and structural complexity. Additional long-read technologies such as Oxford Nanopore ultra-long reads or targeted optical mapping may be required to resolve remaining gaps or collapsed duplications within B-linked regions. Similarly, although Hi-C analysis revealed notable altered chromatin landscape in the 1B genome, these findings are based on single-sample datasets and require further validation. Future studies incorporating biological replicates, cell-type– specific Hi-C, or single-cell 3D genome profiling will be essential to validate and refine these observations, providing a more nuanced understanding of the chromatin effects associated with B chromosome presence.

In conclusion, this study provides a detailed genomic characterization of B chromosome of the important model species, integrating structural, transcriptomic, and 3D architectural dimensions. The high-quality reference genome of *A. latifasciata* offers a valuable resource for future investigations into B chromosome function, chromatin regulation, and genome evolution in cichlids and beyond. Our findings open new avenues for investigating how non-essential, supernumerary chromosomes influence host genome architecture—by mediating structural rearrangements and potentially shaping aspects of genome function. Taken together, our study highlights the complex and dynamic aspects of B chromosomes although non-essential yet influential genomic elements—as potential contributor to shape chromatin reorganization and carrying extra genetic content, raising the possibility that their persistence reflects either functional co-option by the host genome or selfish evolutionary survival.

## Materials and Methods

### Long-read PacBio and Hi-C Genome Sequencing

We obtained 20 adults *Astatotilapia latifasciata* individuals from a local aquarium store in Botucatu, São Paulo, Brazil. Karyotyping was performed at the Integrative Genomics Laboratory (UNESP–São Paulo State University) to determine the presence of B chromosomes. The presence or absence of B chromosomes was further confirmed by genotyping using primers developed by Fantinatti and Martins (2016). High molecular weight genomic DNA was extracted from the muscle tissue of a single adult male carrying one B chromosome (1B) using the Qiagen MagAtrract HMW DNA kit (cat no. 67563), with gDNA extraction for B chromosomes genotyping was performed using phenol-chloroform method following Green and Sambrook (2012).

Sequencing libraries were prepared and sequenced on the PacBio Sequel II platform using SMRT Cell v2.0. For Hi-C sequencing, muscle tissues from one individual with a B chromosome (1B) and one without (0B) were used. Cross-linked chromatin was prepared and libraries constructed by Phase Genomics (Seattle, WA, USA). DNA was digested using the Sau3AI restriction enzyme, and proximity ligation was performed using biotinylated nucleotides to generate chimeric molecules reflecting spatial proximity. These were processed into paired-end sequencing libraries. Chimeric contacts represent loci that are physically proximal in vivo but not necessarily genomically adjacent. All experimental protocols were approved by the Ethics Committee of the Institute of Biosciences, UNESP (Protocol No. 769–2015).

### Chromosome-scale Genome Assembly

Long-read data from the 1B male sample were independently assembled using two tools: Falcon v0.3.0 (Chin et al. 2016) and wtdbg2 v2.5 (Ruan and Li, 2020). Customized parameters were applied for read correction, trimming, overlap detection, and de novo contig/scaffold construction. The assembly generated using wtdbg2 was selected for downstream analysis due to its superior continuity (Table S3).

We polished the assembly using short-read Illumina data previously generated for the same individual (Jehangir et al. 2019) with Pilon v1.22 (Walker et al. 2014). Redundant sequences were removed using Redundans ((Pryszcz and Gabaldón 2016). The curated and polished draft genome was then scaffolded to the chromosome scale using Hi-C data with SALSA2 and Juicer (Durand et al. 2017). Final assembly quality was assessed using gVolante (Nishimura et al. 2017), BUSCO (Simão et al. 2015), and LTR Assembly Index (LAI) (Ou et al. 2018).

### Genome Annotation

Repeat annotation was performed using the Extensive de-novo TE Annotator (EDTA; Ou et al. 2019) and RepeatMasker version 4 using metazoa and crossmatch engine (Smit et al. 2013). Protein-coding genes were predicted using BRAKER (Hoff et al. 2016; Brůna et al. 2021), incorporating transcriptomic evidence mapped by GMAP (Wu et al. 2005) from *A. latifasciata* RNA-seq data. Predicted genes were also compared to the *Metriaclima zebra* genome via BLASTp to identify putative orthologs. BRAKER was run in ab initio mode using GeneMark (Lomsadze et al. 2014) and AUGUSTUS (Stanke et al. 2004). Functional annotation of predicted coding genes on the B chromosome was conducted using BLAST2GO (Conesa et al. 2005), incorporating multiple protein databases for homolog detection.

### B Chromosome Identification

To identify B chromosome–linked sequences, we used male and female short-read Illumina data from 0B, 1B, and 2B individuals. Reads were aligned to the Asta_v3 assembly using Bowtie2 (Langmead et al. 2012) with the “--very-sensitive --end-to-end” alignment mode, and per base coverage was calculated with Bedtools (Quinlan and Hall, 2010). Log2-normalized coverage ratios were compared between 1B and 0B samples to detect B-enriched scaffolds, following the previous adapted strategy by Valente et al. (2014). Multicoverage reads coverage were computed with Bedtools to identify scaffolds enriched in 1B samples.

To further improve B chromosome identification, we analyzed Hi-C contact profiles. Hi-C data from 0B and 1B individuals were aligned to the Asta_v3 genome using HiC-Pro (Servant et al. 2015). Unplaced scaffolds showing significantly higher Hi-C interaction frequencies with the B super-scaffold were identified using Wilcoxon rank-sum tests and incorporated into the B chromosome assembly. Validation of B chromosome sequences was performed by aligning previously published B-specific markers validated by FISH mapping experiments (Valente et al. 2014; Ramos et al. 2017) using BLASTn. Significant hits with high sequence identity (>95%) were considered evidence of correct B chromosome assembly.

### 3D Genome Organization and TAD Analysis

To explore chromatin structure, Hi-C reads from 0B and 1B samples were aligned to the Asta_v3 genome using Juicer (Durand et al. 2017). Differences in 3D conformation were visualized in Juicebox. The in-silico genome was modeled using Chrom3D (Paulsen et al. 2018). The resulting structures were visualized in Chimera (Pettersen et al. 2004). Hi-C matrices were generated and corrected using HiCExplorer v3.0 (Ramírez et al. 2018). Reads were filtered, matrices corrected for bias, and bins with low mapping counts removed. TADs were identified using “hicFindTADs” at 20 kb and 50 kb resolutions with FDR correction. TAD boundaries were visualized using HiCPlotter (Akdemir and Chin 2015). Chromatin loops were detected using “hicDetectLoops” from HiCExplorer, applying a negative binomial model and Wilcoxon test for loop significance. We compared TAD sizes, loop sizes, and duplication breakpoints using Bedtools and regioneR (Gel et al. 2016). Duplication events were inferred using Syri and intersected with TAD boundaries to examine disruption patterns. We also compared TAD profiles of *A. latifasciata* with those of other cichlids (*O. niloticus* and *A. citrinellus*), analyzing Hi-C data with the same pipeline and parameters. Hi-C sequencing data of representative cichlid species including *O. niloticus* and *A. citrinellus* were retrieved from NCBI SRA (Accession: SRR11744829 and SRR12137490) respectively. The Hi-C reads of both representative cichlid genomes data were aligned with their respective assembled chromosome level genome (Assembly NCBI accessions: GCA_013435755.1; GCA_922820385.1). Same set of tools and parameters for Hi-C alignments, matrices correction and prediction of TADs and loops were applied as described for our *A. latifasciata* TADs analysis. We then counted total number of TADs in each species and compared with TADs of 0B and 1B genomes.

### Transcriptome and Gene Expression Analysis

RNA-seq data from brain, muscle, and gonad of 0B and 1B individuals were aligned to the genome using STAR (Dobin et al. 2013). BAM files were sorted and indexed with Samtools (Li et al. 2009). Read counts for B chromosome–localized genes were obtained using Bedtools “multicov”. Log-transformed normalized read counts were analyzed, and ANOVA test was performed to assess significant differences of mean gene expression across chromosomes. Differential expression of B-linked genes was assessed using edgeR v3.36.0 (Robinson et al. 2010), applying CPM > 1, logFC = 1.5, and FDR = 0.05 thresholds. Genes with significant expression changes were identified across tissues.

### Comparative Genomics and Gene Family Analysis

Orthogroup clustering was performed across 11 cichlids and 5 outgroup species using OrthoFinder2 (Emms and Kelly 2019), incorporating DIAMOND (Buchfink et al. 2015) and clustered with OrthoMCL (Li et al. 2003). Phylogenetic trees were reconstructed following multiple sequence alignment with MAFFT (Katoh et al. 2002), alignment trimming using TrimAl (Capella-Gutiérrez et al. 2009), and tree inference with RAxML (Stamatakis 2014) and IQ-TREE (Minh et al. 2020). Divergence times were estimated with MCMCtree (Yang 2000), and convergence assessed in Tracer (Rambaut et al. 2018). Gene family expansions and contractions were assessed with CAFE (Bie et al. 2006). Enrichment analysis of expanded/contracted families was performed in BLAST2GO using Fisher’s exact test (FDR < 0.05).

Structural variants (SVs) between the 0B and 1B genomes were identified using whole-genome alignments generated by NUCmer (Kurtz et al. 2004) and analyzed with SyRI (Goel et al. 2019), a tool designed to detect a broad spectrum of structural rearrangements including inversions, translocations, duplications, and unaligned or divergent regions. Since SyRI requires both the reference and query genomes to be in chromosome-scale assembly format, we first anchored our previously published draft 0B genome (Jehangir et al. 2019) to chromosome scale. To achieve this, we employed a hybrid strategy using Hi-C scaffolding and reference-guided alignment. Specifically, the 0B draft assembly was aligned to the 1B reference genome using RagTag (Alonge et al. 2021), which enables misassembly correction and reference-guided scaffolding based on homology with a high-quality reference. The RagTag-anchored scaffolds were further improved using Hi-C data to resolve ambiguities and finalize chromosome-level continuity (**Table S**). This chromosome-scale 0B genome was then compared with the 1B assembly using SyRI, enabling the detection of genome-wide structural differences, including 1B genome-specific rearrangements potentially associated with B chromosome presence.

## Supplementary Material

Supplementary material including supplementary figures and tables is attached with the manuscript.

## Acknowledgments

Assemblies and all bioinformatics and genomics data analyses were conducted using high-performance computing (HPC) resources at Kasetsart University and UNESP. We thank the members the Integrative Genomics Laboratory at UNESP, and the AGB Research Unit at Kasetsart University for their valuable feedback, early suggestions, and their kind support in this study.

## Authors Contributions

All authors discussed the results and contributed to the final manuscript. M.J. and C.M. conceived the study and, together with S.F.A. and J.I.N.O., designed the experiments and co-wrote the manuscript. M.J. and C.M. secured funding for the project. A.L.C. collected and prepared biological samples. J.I.N.O. extracted DNA and prepared materials for sequencing. M.J. performed genome assembly. S.F.A. and I.R.W. carried out repeat and gene annotation analyses, with support from G.T.V. and M.J. All remaining data analyses and visualizations were conducted by S.F.A. and M.J. K.S. co-supervised the project and provided bioinformatics support for high-performance computing and data processing.

## Data Availability

Sequencing data have been deposited to NCBI’s SRA database under the project accession number PRJNA1283115 with SRA accession IDs SRR34268038, SRR34268037 and SRR34268036. The *A. latifasciata* genome assembly with B chromosome is available under PRJNA1268020. The repeat annotations, gene annotations, and supporting datasets are available on Zenodo (https://doi.org/10.5281/zenodo.15522222).

## Ethics declarations

### Ethics approval and consent to participate

All animal procedures were conducted in compliance with the ethical guidelines established by the Brazilian College of Animal Experimentation. Experimental protocols were reviewed and approved by the institutional ethics committees of the Institute of Biosciences at UNESP (Protocol no. 769/2015) and CEEAAP at UNIOESTE (Protocol no. 13/09).

### Consent for publication

Not applicable.

