## Supplementary figures for "Genome Assembly of *Astatotilapia latifasciata* Uncovers B Chromosome Linked Chromatin Reorganization"

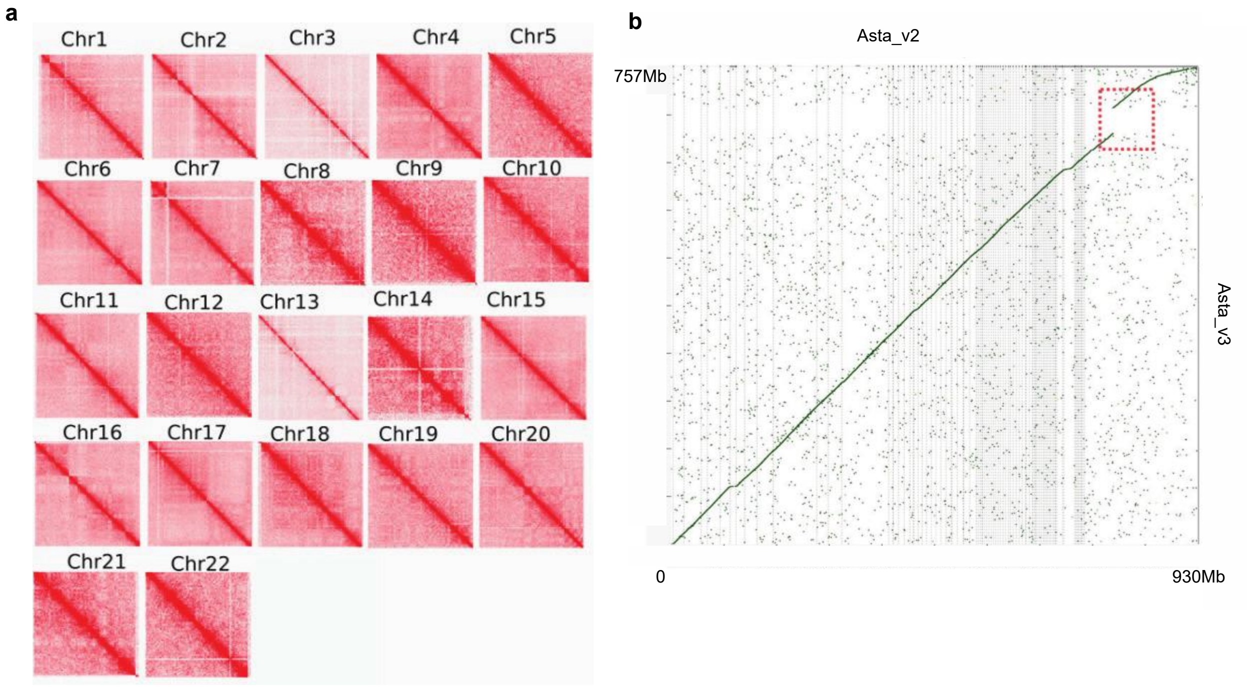


**Fig. S1.** (**a**) Hi-C interaction heatmap at 250 kb bin size resolution for each of the 22 assembled chromosomes of the of genome. (**b**) Comparison of *A. latifascaiata* chromosome level genome (Asta_v3; present study) with the previous *A. latifascaiata* Illumina assembly (Asta_v1; Jehangir et al. 2019) revealing a gap of 150 mb region (red box) that was resolved in upgrading assembly.

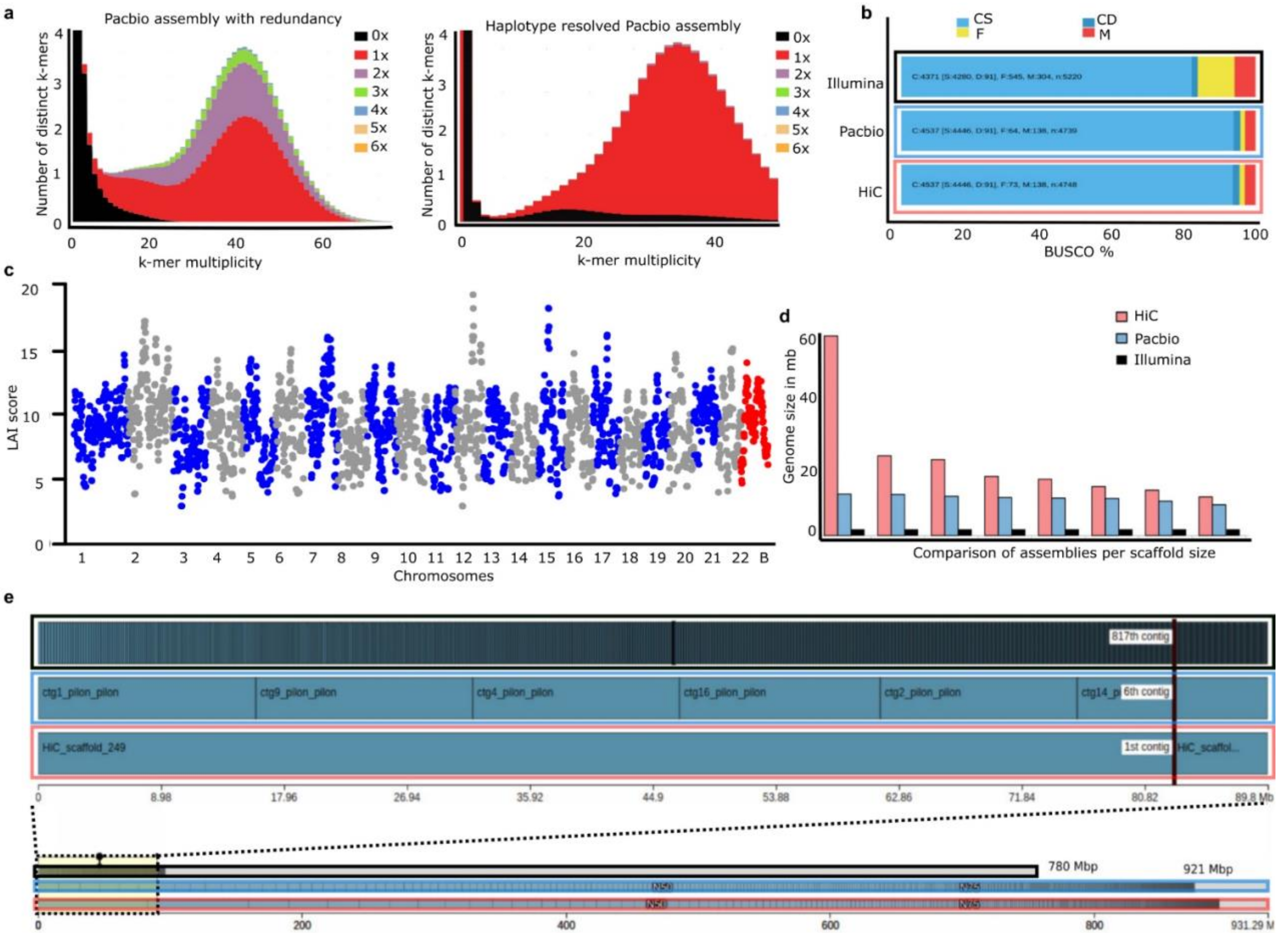


**Fig. S2.** Evaluation of A. latifasciata genome assemblies using multiple quality assessment approaches. **(a)** K-mer spectra plots generated with KAT, showing the distribution of K-mer frequencies in the genome assembly. The left panel displays redundancy in the initial assembly, while the right panel shows improved resolution following haplotype phasing. **(b)** BUSCO analysis comparing genome completeness across assemblies generated using different sequencing technologies. Categories include complete and single-copy genes (light blue), complete and duplicated genes (dark blue), fragmented genes (yellow), and missing genes (red). **(c)** Manhattan plot of the LTR Assembly Index (LAI) scores across the genome, with each dot representing a 3 Mb sliding window adjusted by genome-wide LTR identity, indicating regional assembly quality of repetitive sequences. **(d)** Bar plot comparing scaffold size distributions across assemblies, illustrating improvements in contiguity from kilobase to megabase scale. **(e)** Icarus visualization from QUAST

(Mikheenko et al., 2016) displaying the improved continuity in the Hi-C–scaffolded assembly. An example chromosome is shown, highlighting its formation from multiple contigs derived from Illumina and PacBio datasets. Each vertical bar denotes a contig or scaffold, with box sizes proportional to their genomic length.

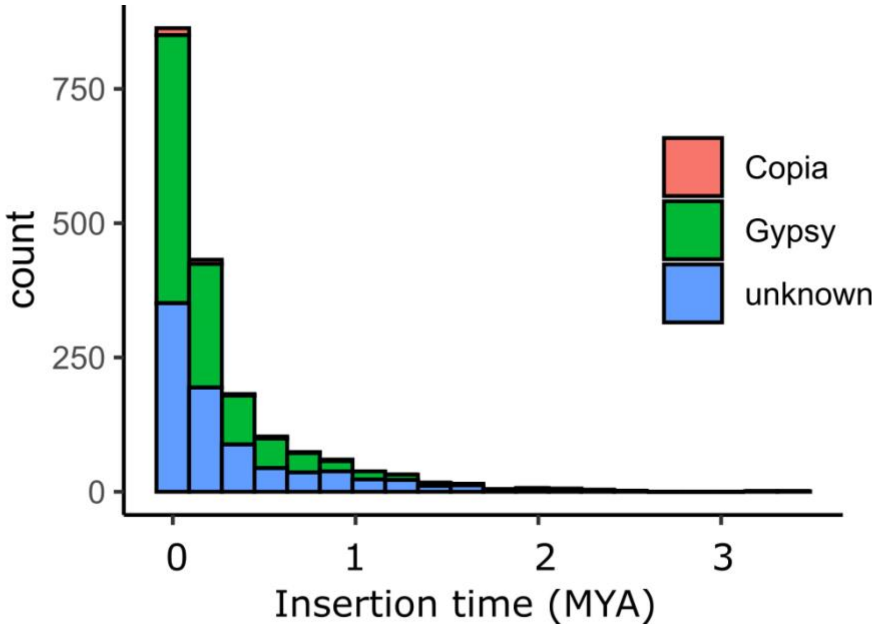

**Fig. S3.** Insertion age of LTR elements in millions of years (Mya). Comparison of TEs (Copia, Gypsy and unknown) insertion time in the last 3 Mya for major groups of LTR retrotransposons.

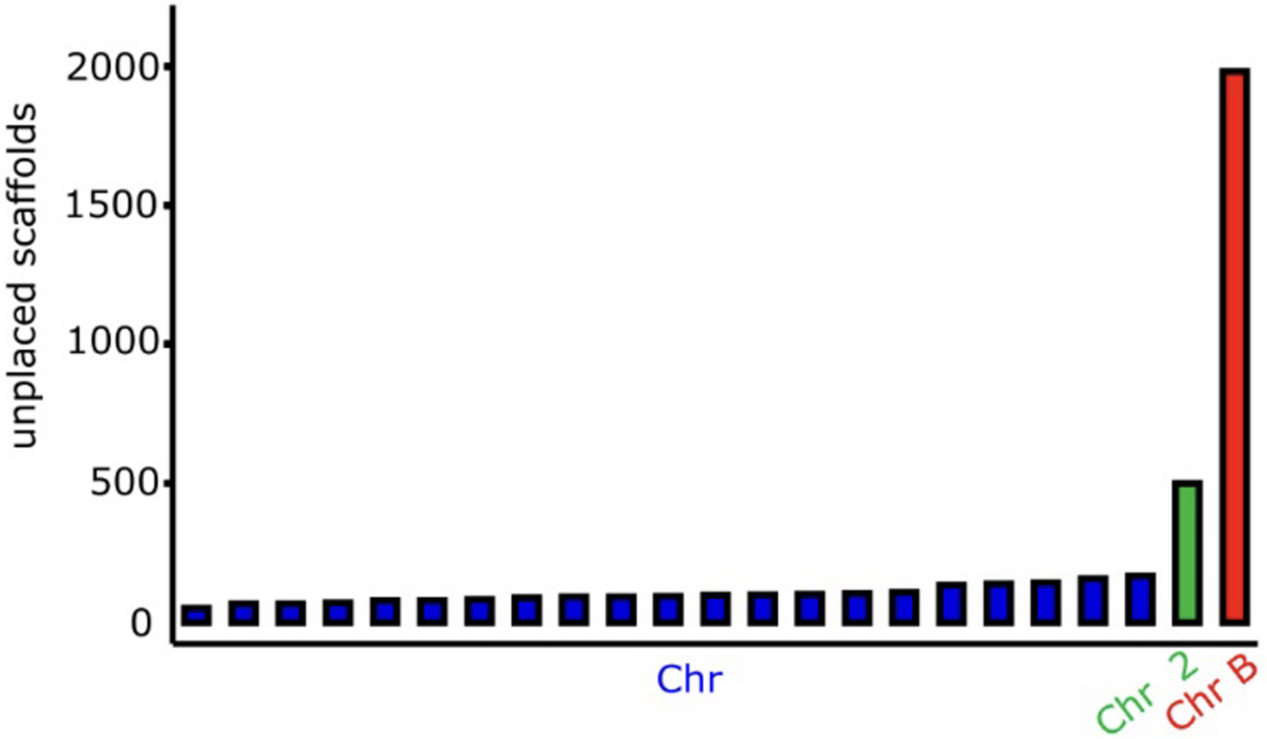


**Fig. S4.** The bar chart shows the similarity mapping indicating the association of unplaced scaffolds with assembled chromosomes. The blue bars in the chart represent A chromosomes, except Chr 2 and Chr B shown as green and red bars respectively. A substantial number of unplaced scaffolds shown association (the highest number of paralogs) with Chr B.

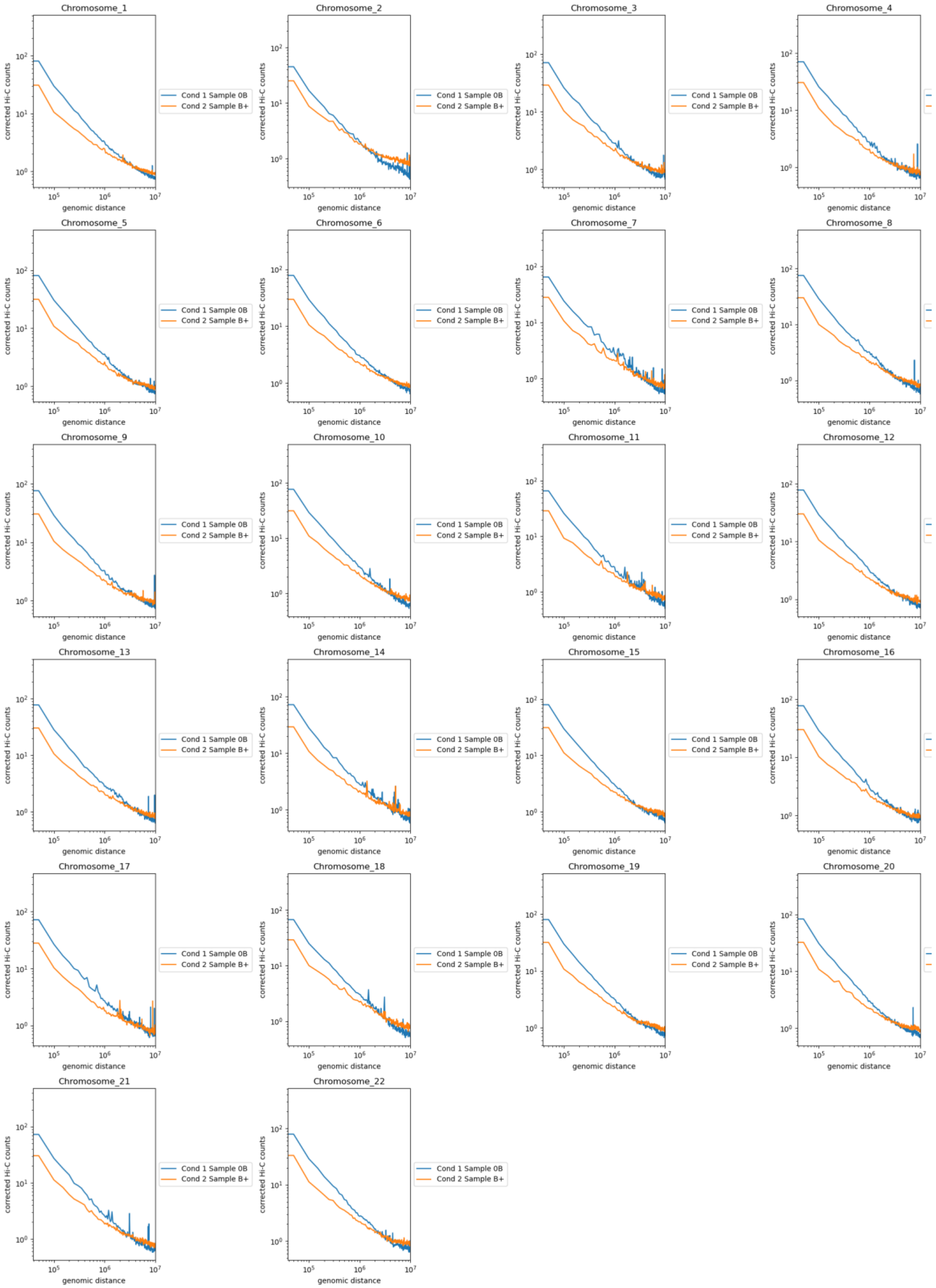


**Figure S5.** Line plots show the chromosome-wise distribution of contact probability ($P_S$) which was calculated as a function of genomic distance for interactions within individual chromosome arms across the whole genome of 0B and 1B.

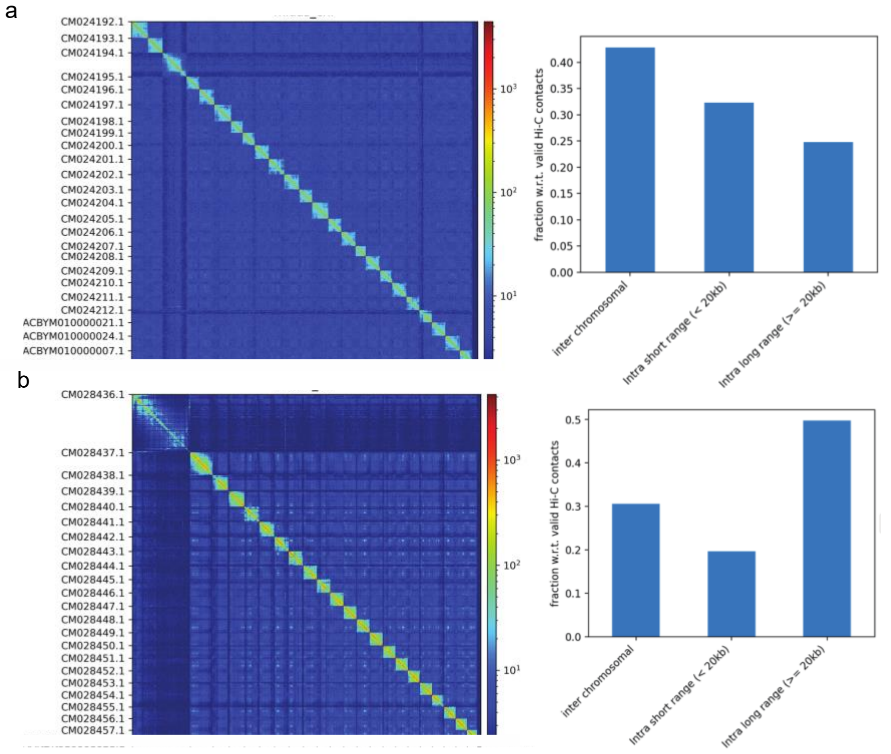


**Figure S6.** Hi-C heatmap based on the chromosome-scale assembly of the *A. citrinellus* **(a)** and *O. niloticus* **(b)** genomes. The heatmaps represent the contact matrices generated by aligning the Hi-C data to the chromosome-scale assemblies of the *A. citrinellus* and *O. niloticus* and bar graphs represent the different interaction types.

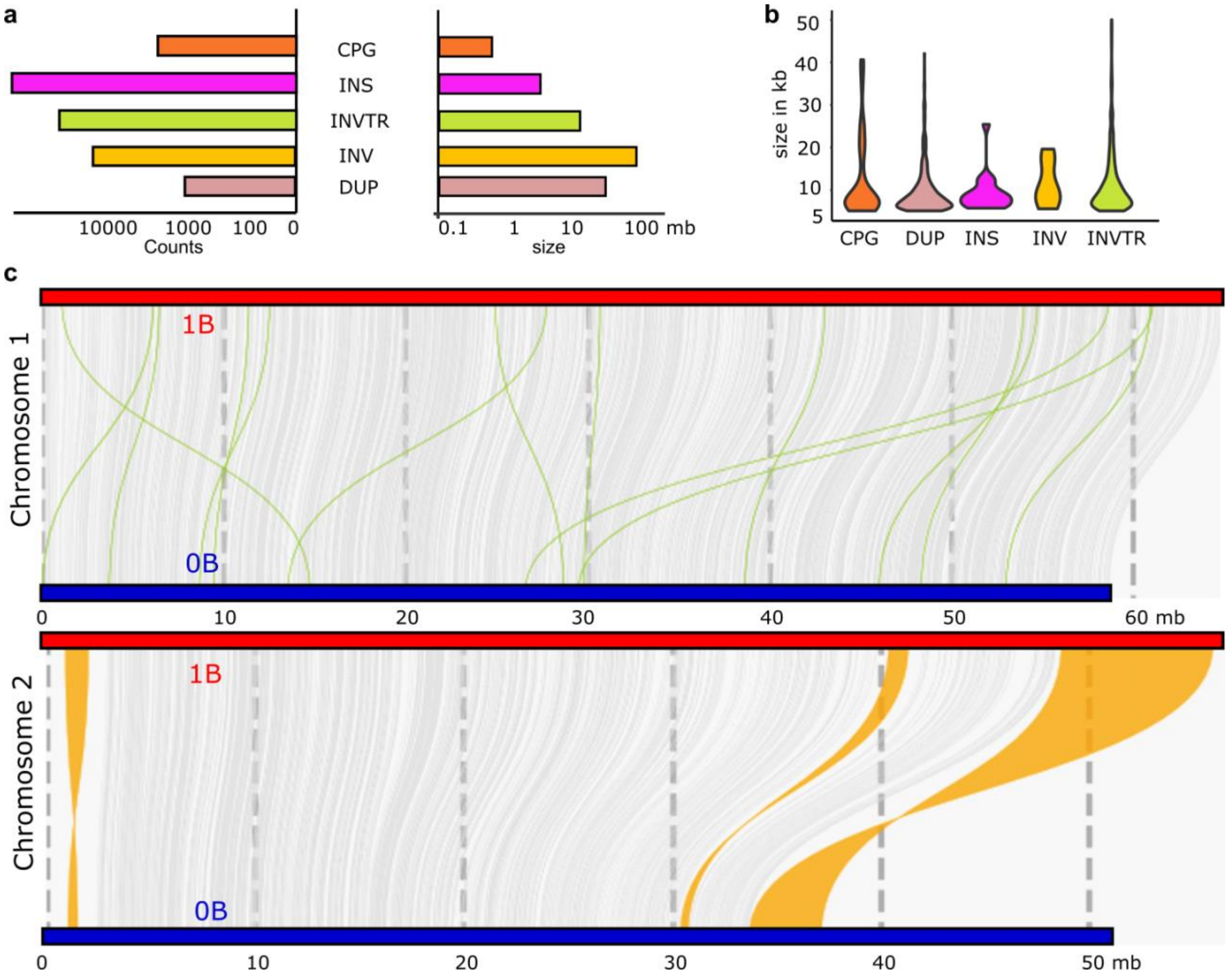


**Figure S7**. Genomic differences between the 0B and 1B genomes and associated patterns of rearrangements. (**a)** The bar graphs show the total number of different rearrangements and their total length in 1B. (**b)** The violin plot depicts the length distribution of each rearranged region in 1B genome. The distribution of rearrangements with at least 5 Kb length is plotted. Refer to Fig. S13 for distribution of smaller size rearrangements. **c** Example of intra-chromosomal rearrangements (translocations, inversions and syntenic shown as green, orange and grey color respectively) between 0B and 1B genomes

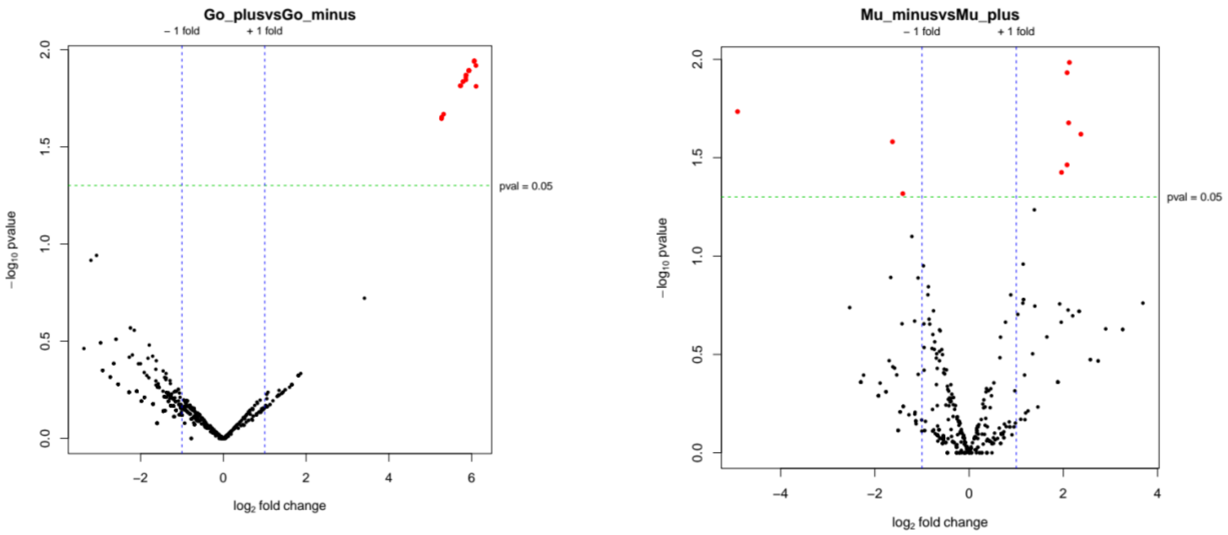


**Figure S8.** Comparison of DEGs between the gonads and muscles tissues of 0B and 1B genomes. Scatter plot showing the correlation of gene abundance. DEGs are shown as red dots and black dots indicate genes that lack of significantly difference.

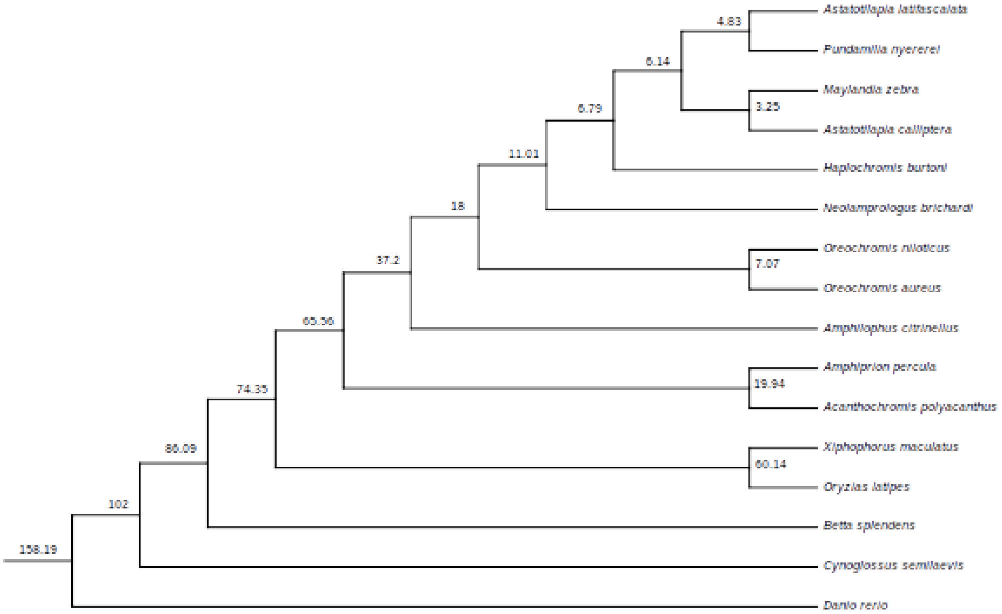


**Fig S9.** Phylogenetic tree and estimated divergence times of cichlid species. A maximum-likelihood phylogenetic tree was inferred from concatenated protein sequences corresponding to 364 shared single-copy genes. Species divergence was estimated using a Bayesian relaxed molecular clock model.

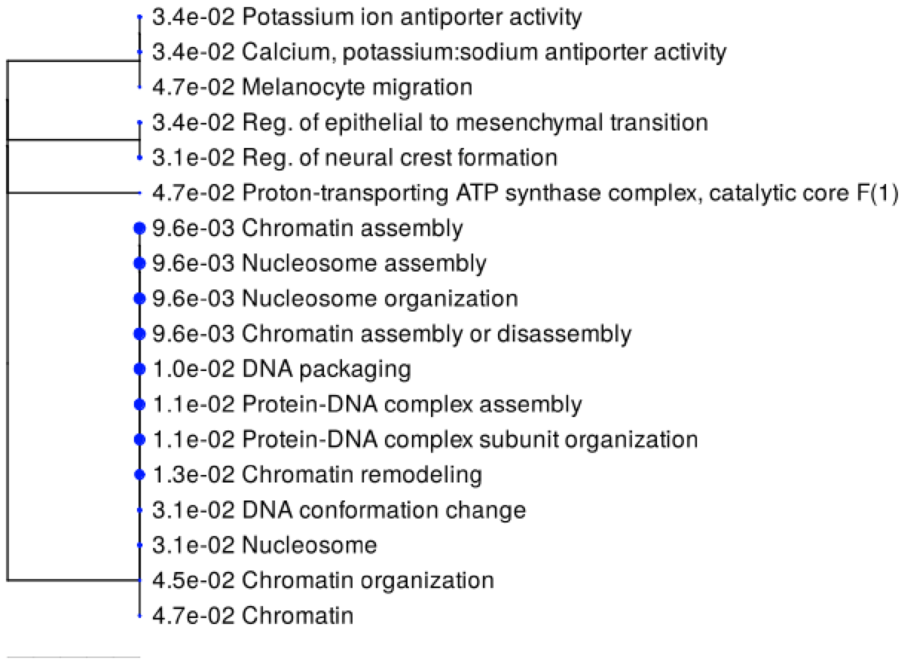


**Figure S10.** Graphical representation of summarized tree of the enriched gene ontology (GO) terms of CAFE expanded genes on B chromosome.

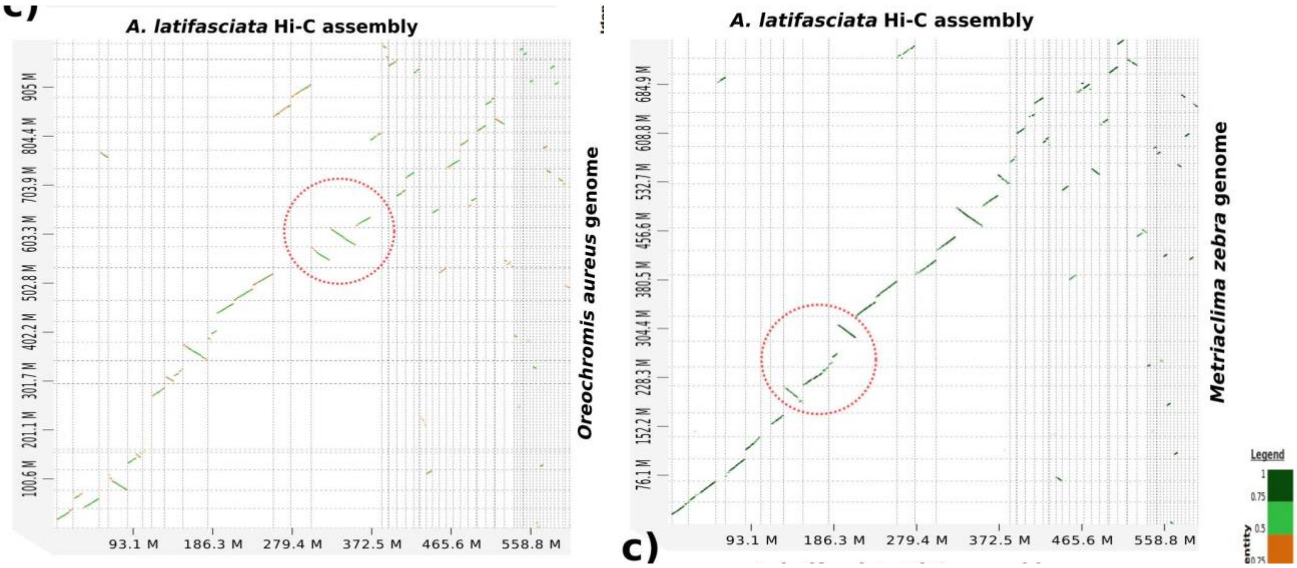


**Figure S11**. Comparative genomic sytenic analysis of *M. zebra* and *A. aureu*s genomes with *A. latifascaiata* genome. The x-axis represents one organism's genome, and the y-axis represents the other. Each box within the graph represents one chromosome/scaffold. These genomes are compared against one another and every region in one genome that matches respective region in another organism is indicated by a dot which increases with alignment size with varying color based on alignment score. Jumps in green lines signify an insertion or deletion in one of the genomes and inverted lines show inversion, whereas a line in the same box of one genome against the multiple lines of other genome corresponds to fusion. Note that the alignment gap (highlighted dotted red box) in the panel (a) represents a false deletion because of the missing region in the Illumina assembly. Red dotted circles show the cross-species chromosomal rearrangements including inversion and fusion.
